# Midkine noncanonically suppresses AMPK activation through disrupting the LKB1-STRAD-Mo25 complex

**DOI:** 10.1101/2021.09.27.462083

**Authors:** Tian Xia, Di Chen, Xiaolong Liu, Huan Qi, Wen Wang, Huan Chen, Ting Ling, Wuxiyar Otkur, Chen-Song Zhang, Jongchan Kim, Sheng-Cai Lin, Hai-long Piao

**Affiliations:** CAS Key Laboratory of Separation Science for Analytical Chemistry, Dalian Institute of Chemical Physics, Chinese Academy of Sciences, Dalian 116023, China.; State Key Laboratory for Cellular Stress Biology, School of Life Sciences, Xiamen University, 361102 Fujian, China.; Department of Life Sciences, Sogang University, Seoul 04107, Republic of Korea; University of Chinese Academy of Sciences, Beijing 100049, China.

## Abstract

Midkine (MDK), an extracellular growth factor, regulates signal transduction and cancer progression by interacting with receptors, and it can be internalized into the cytoplasm by endocytosis. However, its intracellular function and signaling regulation remain unclear. Here, we show that intracellular MDK interacts with LKB1 and STRAD to disrupt the LKB1-STRAD-Mo25 complex. Consequently, MDK decreases the activity of LKB1 to dampen both the basal and stress-induced activation of AMPK by glucose starvation or treatment of 2-DG. We also found that MDK accelerates cancer cell proliferation by inhibiting the activation of the LKB1-AMPK axis. In human cancers, compared to other well-known growth factors, MDK expression is most significantly upregulated in cancers, especially in liver, kidney and breast cancers, correlating with clinical outcomes and inversely correlating with *PRKAA1* (encoding AMPKα1) expression and phosphorylated AMPK levels. Our study elucidates an inhibitory mechanism for AMPK activation, which is mediated by the intracellular MDK through disrupting the LKB1-STRAD-Mo25 complex.

## INTRODUCTION

AMP-activated protein kinase (AMPK), consisting of catalytic subunit α and regulatory subunit β and γ (Lin & Hardie, 2018), is the core cellular energy sensor and regulator (Garcia & Shaw, 2017). Under energy stress conditions, an elevated cellular AMP/ATP ratio induces conformational changes in the AMPK heterotrimer and induces the exposure of the AMPKα Thr172 site (Hardie, 2018, Oakhill, Steel et al., 2011), which can be phosphorylated by upstream kinases (Carling, 2017). Phosphorylation at Thr172 leads to the activation of AMPK (Hawley, Davison et al., 1996, Suter, Riek et al., 2006), which directly phosphorylates a series of substrates to postpone energy-consuming processes, such as cell proliferation and fatty acid synthesis, and to promote energy-producing procedures, including catabolism and autophagy (Goodman, Liu et al., 2014, Mihaylova & Shaw, 2011). AMPK is closely related to diverse diseases (Rider, 2016), including dual and controversial roles in cancer (Faubert, Vincent et al., 2015, Jeon & Hay, 2015, Russell & Hardie, 2020). Although defined as a tumor suppressor by many studies (Faubert, Boily et al., 2013, Houde, Donzelli et al., 2017, Huang, Wullschleger et al., 2008, Vara-Ciruelos, Dandapani et al., 2019), AMPK promotes cancer progression under certain conditions by rescuing cancer cells from nutrient deficiency (Eichner, Brun et al., 2019, Laderoute, Calaoagan et al., 2014, Saito, Chapple et al., 2015, Shaw, 2015). LKB1, CAMKKβ and TAK1 are upstream kinases of AMPK that all phosphorylate AMPKα at the Thr172 site (Goodman et al., 2014). Among these proteins, the serine/threonine kinase LKB1 mediates the best-characterized classical AMPK activation route(Woods, Johnstone et al., 2003), especially in cancer cells. In contrast to most kinases, which are usually phosphorylated and activated by upstream kinases, LKB1 forms a heterotrimer with pseudokinase STRAD and scaffolding protein Mo25 and then undergoes self-phosphorylation at multiple amino acids to self-induce its kinase activity (Hawley, Boudeau et al., 2003, Zeqiraj, Filippi et al., 2009a, Zeqiraj, Filippi et al., 2009b). Some studies have demonstrated that disruption of the LKB1-STRAD-Mo25 complex decreases AMPKα Thr172 phosphorylation levels and attenuates AMPK activity (Lin, Elf et al., 2015).

Midkine (MDK, encoded by the *MDK* gene) is a pleiotrophin family growth factor that plays vital roles in different physiological processes, such as embryo and nerve development, blood pressure control, inflammation and immune response (Kadomatsu, Bencsik et al., 2014, Muramatsu, 2010, Sorrelle, Dominguez et al., 2017, Yoshida, Sakakima et al., 2014). MDK is highly expressed in different types of cancer (Kato, Shinozawa et al., 2000, Meng, Tan et al., 2015, Shaheen, Abdel-Mageed et al., 2015) and promotes tumor progression by positively regulating cell proliferation, invasion and migration (Rawnaq, Dietrich et al., 2014, Sun, Hu et al., 2017, Xu, Qu et al., 2009, Yao, Li et al., 2014). The molecular weight of mature MDK is 13 kD after the cleavage of the signal peptide (Muramatsu, 2014). As a secreted protein, MDK binds transmembrane receptors (Kadomatsu, Kishida et al., 2013), including PTPξ, ALK, Notch2 and LRP1, and thus activates intracellular signaling (Herradon, Ramos-Alvarez et al., 2019, Kadomatsu et al., 2013, Kishida, Mu et al., 2013, Lorente, Torres et al., 2011, Muramatsu, Zou et al., 2000). It has also been reported that extracellular MDK can be transported into the cytosol by endocytosis and then enter the nuclei where it undergoes proteasomal degradation, but the intracellular functions of MDK are still unclear (Dai, Shao et al., 2008, Shibata, Muramatsu et al., 2002, Suzuki, Shibata et al., 2004).

Here, we report that MDK suppresses AMPK activation in the cytoplasm instead of acting as an extracellular ligand. In the cytosol, MDK interacts with LKB1 and STRAD to depolymerize the LKB1-STRAD-Mo25 complex and reduce LKB1 activity, consequently reducing the phosphorylation of AMPKα. Decreasing the cellular MDK expression level or maintaining the extracellular localization of MDK elevates AMPKα phosphorylation in cells. In cancer cells, MDK promotes cell proliferation by suppressing the LKB1-AMPK axis, and MDK expression correlates with clinical outcomes and inversely correlates with LKB1/AMPK signaling pathway activation. Therefore, our study reveals a previously undescribed molecular function and mechanism of MDK, which may facilitate further clinical application of MDK in targeted cancer therapy.

## RESULTS

### Midkine suppresses AMPK activation in an intracellular localization-dependent manner

Most previous MDK studies focused on identifying the transmembrane receptors of MDK to connect the secreted MDK with intracellular signaling. Very few studies have reported that extracellular MDK can be internalized into cells by endocytosis and localize near the nucleus to regulate rRNA synthesis (Dai, 2009, Dai et al., 2008). To confirm this localization, we examined the transport and relocalization of MDK. Consistent with previous studies, MDK was secreted into the cell medium of HepG2 and HCCLM3 cells expressing high levels of MDK (Fig 1A), and this secreted MDK was internalized into Bel-7402, SMMC-7721 and MHCC97H cells not expressing MDK (Figs 1B and EV1A and B). This internalized intracellular MDK was found 15 minutes after MDK-overexpressing cell culture medium (or conditioned medium, CM) treatment (Figs EV1A and B), suggesting that the intracellular relocalization of MDK was efficient in MDK non-expressed cells. However, intracellular MDK was mostly localized in the cytoplasm, not in the nucleus (Figs 1C and EV1C and D). This phenomenon indicated that MDK may possess an unexplored function in the cytoplasm.

**Figure 1.**
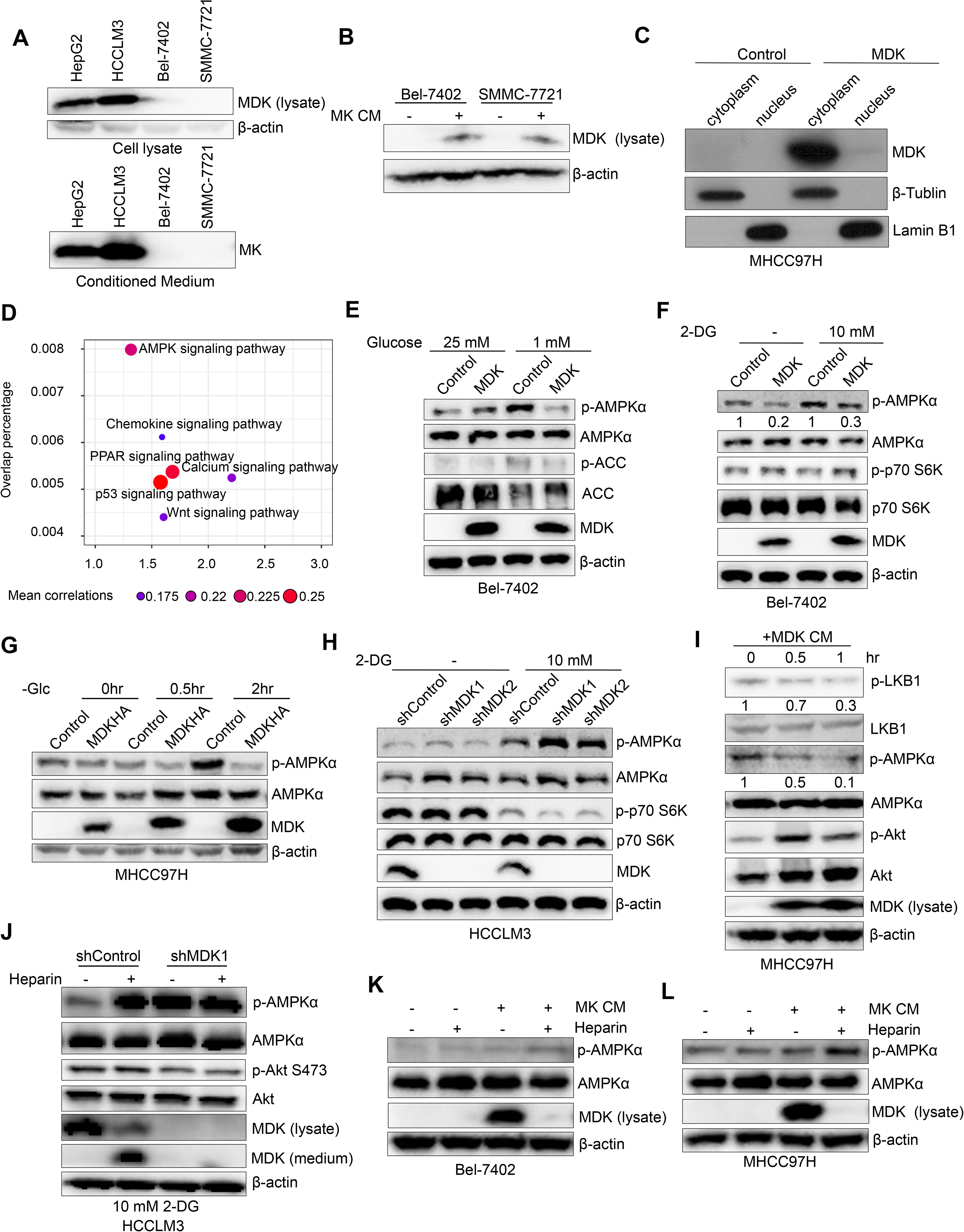
Midkine suppresses AMPK activation in an intracellular localization-dependent manner. **A.** Western blotting of MDK and β-actin in the HepG2, HCCLM3, Bel-7402 and SMMC-7721 cell lysate and conditioned medium (CM). **B.** Western blotting of MDK and β-actin in the Bel-7402 and SMMC-7721 cells with or without CM from MDK-overexpressing MHCC97H cells. **C.** Western blotting of MDK, β-tubulin and Lamin B1 in the cytoplasmic and nuclear fractions of the MHCC97H cells transduced with MDK and the control cells. **D.** Signaling pathway enrichment assay with MDK-correlated genes based on the TCGA database. **E.** Western blotting of the Bel-7402 cells transduced with MDK and the control cells treated with different concentrations of glucose (25 mM and 1 mM) for 2 hours. **F.** Western blotting of the Bel-7402 cells transduced with MDK and the control cells with or without 10 mM 2DG treatment for 4 hours. **G.** Western blotting of the MHCC97 cells transduced with MDK and the control cells at different times during glucose starvation. **H.** Western blotting of the HCCLM3 cells transduced with two independent MDK shRNAs with or without 10 mM 2DG treatment for 4 hours. **I.** Western blotting of the MHCC97H cells at different time points of CM treatment. CM is from MDK-overexpressing MHCC97H cells. **J.** Western blotting of the HCCLM3 cells transduced with MDK shRNA1 and the control cells with or without heparin treatment (30 μg/ml) for 4 hours under 10 mM 2DG culture conditions. **K and L.** Western blotting of the Bel-7402 (K) and MHCC97H (L) cells with or without combined treatment with MDK-overexpression CM and heparin (30 μg/ml).

To identify the MDK-regulated cell signaling pathways, we first performed a pathway enrichment analysis. Interestingly, we found that the AMPK signaling pathway was the most highly correlated with MDK among the pathways (Fig 1D). To further examine the relationship between MDK and the AMPK pathway, we tested the level of phosphorylated AMPKα at Thr172 in MDK-knockdown and MDK-overexpressing cells. In both Bel-7402 and MHCC97H cells, the overexpression of MDK inhibited the level of phosphorylated AMPKα during glucose starvation, 2-DG stimulation or FBS deprivation (Figs 1E, F and G, and EV1E). In contrast, knocking down MDK expression by shRNA led to elevated levels of AMPKα phosphorylation (Figs 1H and EV1F and G), and restoring MDK expression decreased AMPKα phosphorylation (Fig EV1F and G). Taken together, these results suggest that MDK suppresses AMPK activation in human cancer cells.

MDK is well known to be secreted after posttranslational modification. Although a number of studies have reported that extracellular MDK can be transported into cells, the function of intracellular MDK remains unclear. Therefore, to understand whether internalized cellular MDK is critical for AMPK repression, we investigated the effect of extracellular MDK on AMPK activation. We collected CM containing secreted MDK from MDK-overexpressing MHCC97H cells and then used it to culture MDK-deficient MHCC97H parental cells. Upon the application of CM from MDK-overexpressing cells, the intracellular MDK level increased in a time-dependent manner, and this treatment, which triggered AKT phosphorylation as previously reported(Sandra, Harada et al., 2004), decreased LKB1 and AMPKα phosphorylation (Fig 1I). In addition, Bel-7402 cells cultured with control CM exhibited increased AMPK activation after 2-DG treatment, however, the cells cultured with CM from MDK-overexpressing showed decreased AMPK activation, even after 2-DG treatment (Fig EV1H).

Next, to further clarify whether the intracellular relocalization of MDK is indispensable for AMPK suppression, we induced the transportation of MDK into the cytoplasm. MDK is a heparin-binding protein(Iwasaki, Nagata et al., 1997, Kadomatsu et al., 2013). Heparin is a sulfated glycosaminoglycan polymer that can bind MDK and restrict its movement to the inside of cells (Kishida & Kadomatsu, 2014, Muramatsu, Yokoi et al., 2011). By adding heparin to the medium of the HCCLM3 cells, the intracellular MDK level decreased dramatically, and MDK significantly accumulated in the cell culture medium (Fig 1J). Heparin reduced intracellular MDK and obviously elevated AMPKα phosphorylation in the HCCLM3 cells (Fig 1J). In contrast, knocking down MDK did not alter AMPKα phosphorylation levels in cells during heparin application (Fig 1J), excluding the possibility that AMPK is activated by heparin. Similarly, heparin decreased the intracellular MDK level in the MDK-overexpressing cells and promoted AMPKα phosphorylation but did not alter the AMPK activity in the Bel-7402 control cells (Fig EV1I). Additionally, heparin caused decreased intracellular MDK levels and elevated AMPKα phosphorylation in MDK-CM-treated Bel-7402 and MHCC97H cells (Fig 1K and L). In summary, these results indicate that intracellular MDK suppresses AMPK phosphorylation in a cytosol-dependent manner and corroborates findings indicating an intracellular function for MDK.

### Midkine associates with AMPK subunits and its upstream regulating factors

Since MDK regulates the phosphorylation of LKB1 and AMPK, we tried to reveal the underlying mechanisms by which MDK functions. First, we isolated MDK-associated protein complexes in HEK293T cells through tandem affinity purification followed by mass spectrometry (MS) analysis. According to their biological functions, the MDK-associated proteins were classified into different groups (Fig EV2A). We found that some MDK-associated proteins belonged to LKB1 substrates, AMPK regulators or metabolic regulation factors (Figs 2A and EV2A). Interestingly, MDK associated with the LKB1 substrates MARK and SIK3 of AMPK family proteins, and the well-studied AMPK ubiquitination regulators USP10 and UBE2O were also added to the prey list (Fig 2A).

**Figure 2.**
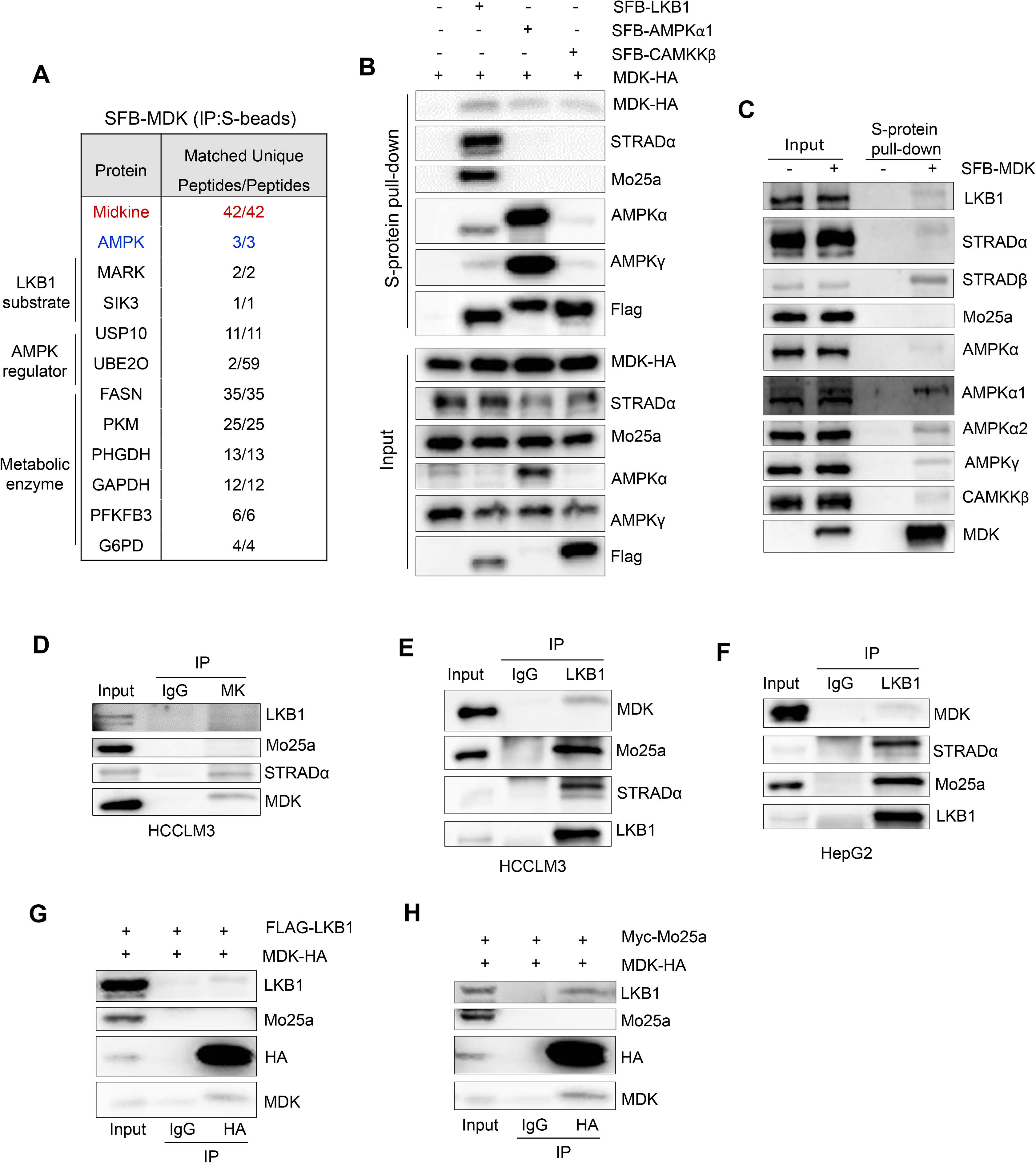
Midkine associates with AMPK subunits and its upstream regulating factors. **A.** AMPK kinase subunit and LKB1 substrates were identified as MDK-associated proteins in HEK293A cells by tandem affinity purification–mass spectrometry. Bait MDK protein is marked in red. AMPKα is marked in blue. The right column of numbers represents unique peptide number/total peptide number. **B and C.** MDK associates with LKB1, CAMKKβ and AMPK subunits. The indicated constructs were expressed in the HEK293T cells for 24 hours, and cell lysates were subjected to pull-down assays with S protein beads. **D.** MDK was immunoprecipitated from HCCLM3 cells and subjected to Western blot analysis with antibodies against LKB1, Mo25a, STRADα and MDK. **E and F.** LKB1 was immunoprecipitated from HCCLM3 (E) and HepG2 (F) cells and subjected to Western blot analysis with antibodies against MDK, Mo25a, STRADα and LKB1. **G and H.** HEK293T cells were cotransfected with MDK-HA and Flag-tagged LKB1 (G) or Myc-tagged Mo25a (H) and coimmunoprecipitated with HA primary antibody and subjected to Western blot analysis with antibodies against LKB1, HA, Mo25a and MDK.

The specific interaction between MDK and the AMPK α subunit, as well as the well-studied AMPK upstream kinases LKB1 and CAMKKβ, was confirmed by pull-down assays (Figs 2B and EV2B and C). Additionally, MDK form complex with endogenous AMPK signaling components such as LKB1, STRADα/β, AMPKα1/2, AMPKγ and CAMKKβ in MDK-transduced HEK293T cells (Fig 2C). Furthermore, coimmunoprecipitation (co-IP) assays showed that LKB1 can be detected in endogenous MDK immunoprecipitates from HCCLM3 cells (Fig 2D) and that endogenous MDK is pulled down with endogenous LKB1 immunoprecipitates from HCCLM3 and HepG2 cells (Fig 2E and F). Interestingly, we noticed that STRADα and Mo25a, which bind LKB1 and facilitate LKB1 activation, showed different interaction ability with MDK (Figs 2D-H and EV2B and C). STRADα was associated with both endogenous and exogenous MDK; however, Mo25a was not detected in the either the endogenous co-IP or exogenous pull-down assays (Figs 2C, D, G and H, and EV2B and D). These results suggested that MDK may inhibit the protein machinery of LKB1-Mo25-STRAD.

Considering that LKB1 phosphorylates AMPKα at Thr172, we wondered whether MDK interacts with LKB1 and AMPKα directly or indirectly via the kinase-substrate reaction. To test this hypothesis, we stably expressed MDK in LKB1-deficient A549 cells and found that AMPKα was detected only in the MDK coprecipitates from LKB1-reconstituted A549 cells ( Fig EV2D). This result indicated that the interaction between MDK and AMPK may be dependent on LKB1. LKB1 contains a kinase domain in the middle of its amino acid sequence (Fig EV2E). Although AMPKα was associated with different forms of LKB1 (the full length protein, the N-terminus with the kinase domain only and the C-terminus only), MDK interacted only with the NK domain (amino acids 1-309), which was similar to STRAD and Mo25 (Fig EV2E and F). In summary, we discovered that both MDK and AMPKα are physically associated with LKB1 through its N-terminal kinase domains in cells.

### Midkine suppresses AMPKα activation through interacting with LKB1

LKB1 and calcium/calmodulin-dependent protein kinase kinase β (CAMKKβ) are well-known AMPK upstream kinases, and both kinases can phosphorylate the AMPK α subunit at the Thr172 site(Fogarty, Ross et al., 2016, Woods et al., 2003). Although MDK associates with LKB1 and CAMKKβ (Fig 2B and C), whether MDK regulates AMPKα activities through LKB1 or CAMKKβ remains unclear. Interestingly, the activity of LKB1 and AMPKα was increased in MDK-knockdown Hep3B and HCCLM3 cells (Fig 3A and B). In contrast, restoring MDK expression in these knockdown cells recovered LKB1 activity and AMPKα phosphorylation (Fig 3A and B), suggesting that MDK contributed to the LKB1-modulated AMPKα suppression. Furthermore, in LKB1-deficient A549 cells, MDK overexpression did not alter AMPKα phosphorylation levels until LKB1 expression was restored (Fig 3C and D), indicating that MDK suppressed AMPKα activation through LKB1. In HCCLM3 cells, AMPKα phosphorylation was elevated when the intracellular MDK levels were decreased by heparin treatment, but the effect of heparin was attenuated in LKB1-knockdown cells (Fig 3E). These results indicated that LKB1 is involved in the MDK-induced regulation of AMPK activity.

**Figure 3.**
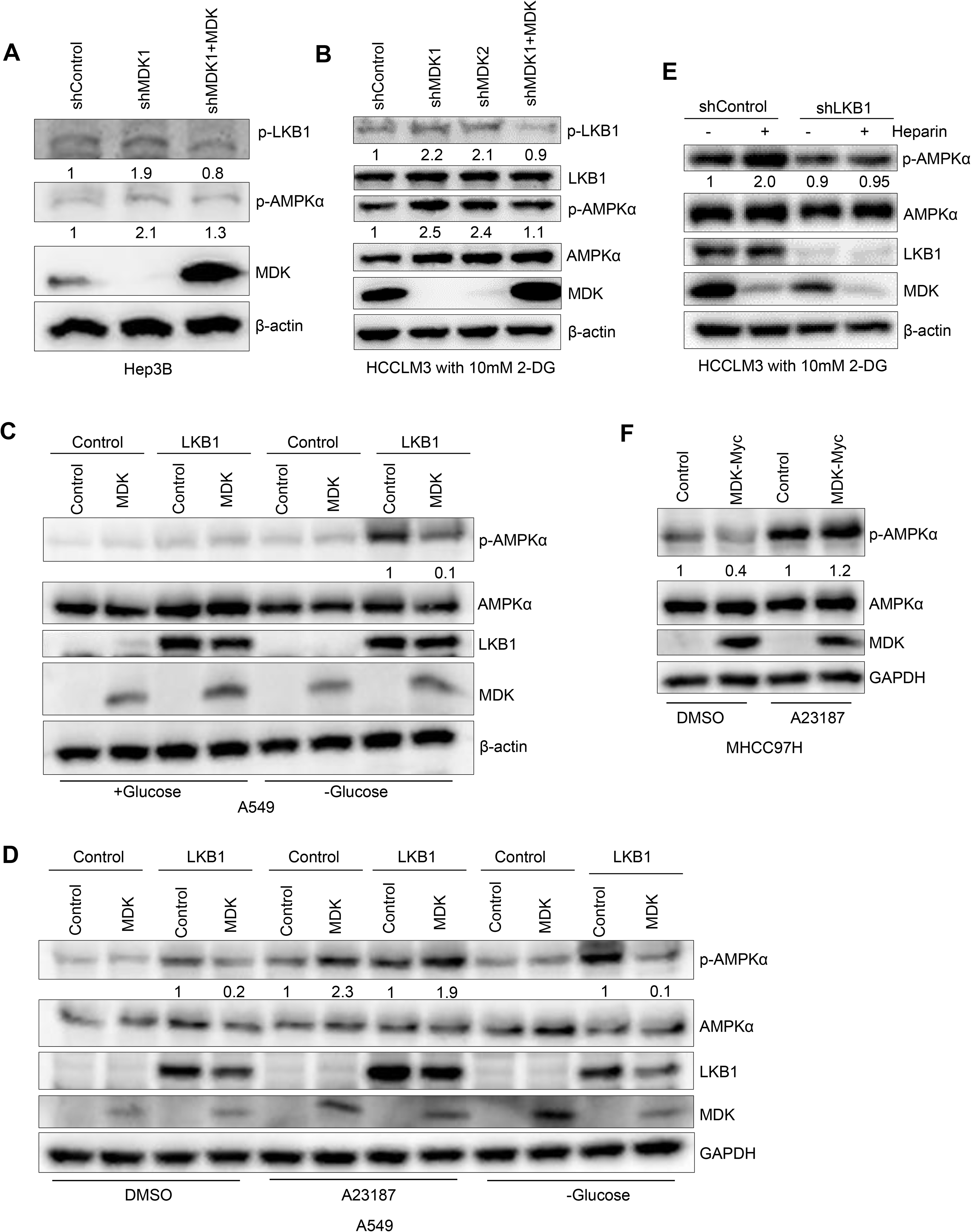
Midkine suppresses AMPKα activation through LKB1. **A and B.** Western blot analysis of the Hep3B cells transduced with MDK shRNA1 and restored MDK in the MDK-knockdown cells (A) and the HCCLM3 cells transduced with two independent MDK shRNAs and restored MDK in the MDK-knockdown cells after 4 hours of 10 mM 2-DG treatment (B). Western blot analysis was performed with antibodies against p-LKB1, LKB1, p-AMPKα, AMPKα, MDK and β-actin. **C and D.** Western blot analysis of the A549 cells transduced with LKB1 alone or in combination with MDK and the control cells with glucose starvation for 2 hours (C) and the A549 cells transduced with LKB1 alone or in combination with MDK and the control cells with DMSO or A23187 (10 μg/ml) treatment or glucose starvation for 2 hours (D). Western blot analysis was performed with antibodies against p-AMPKα, AMPKα, LKB1, MDK and β-actin. **E.** Western blot analysis of p-AMPKα, AMPKα, LKB1, MDK and β-actin in the HCCLM3 cells transduced with LKB1 shRNA and the control cells with or without heparin (30 μg/ml) treatment and 10 mM 2-DG treatment for 4 hours. **F.** Western blot analysis of p-AMPKα, AMPKα, MDK and β-actin from the MHCC97H cells transduced with MDK-Myc and the control cells treated with DMSO or A23187 (10 μg/ml).

To understand whether MDK mediates AMPK signaling through CAMKKβ, we used CAMKKβ activator A23187 to treat MDK-restoring MHCC97H cells. The overexpression of MDK suppressed AMPK activation upon DMSO treatment; however, AMPKα phosphorylation was elevated regardless of the MDK expression after CAMKKβ activator A23187 treatment (Fig 3F). In agreement with this, A23187 stimulated AMPKα activation regardless of the level of MDK or LKB1 expression, while MDK suppressed AMPKα phosphorylation in the presence of LKB1 during glucose starvation (Fig 3C and D). Considering these results, we speculated that MDK mediates AMPK activity through LKB1.

### Midkine disrupts LKB1-STRAD-Mo25 complex

LKB1, a serine/threonine kinase, forms a heterotrimeric complex with pseudokinase STRAD and scaffolding-like adaptor Mo25a and undergoes conformational change and self-phosphorylation to achieve full activation (Zeqiraj et al., 2009a). Our results demonstrated that MDK-mediated repression of AMPK activation relied on LKB1 (Fig 3C-E), and the level of MDK expression was correlated with LKB1 phosphorylation (Fig 3A and B). In addition, we found that MDK physically interacts with LKB1 and STRAD (Fig 1B-H). Considering that the formation of the LKB1-STRAD-Mo25a complex necessary for LKB1 activity, we surmised that MDK affects the stability of the LKB1-STRAD-Mo25 heterotrimer.

To investigate this hypothesis, we performed coimmunoprecipitation assays with LKB1 from MDK-transduced cells. The overexpression of MDK significantly inhibited the formation of the LKB1-STRAD-Mo25a complex in endogenous LKB1 immunoprecipitates from HEK293T cells (Fig 4A). In contrast, knocking down MDK increased the level of STRAD and Mo25 in the endogenous LKB1-containing immunoprecipitates from HCCLM3 cells (Fig 4B). Furthermore, to prevent the phosphorylation of LKB1 from affecting this interaction, we expressed FLAG-tagged wild-type (WT) and kinase dead (KD, K78L) LKB1 in HEK293T cells. As previously reported, both the WT and KD LKB1 bound strongly to STRAD and Mo25a (Fig EV3A). Next, we simultaneously expressed FLAG-tagged LKB1-KD and SFB-tagged AMPKα1 in HEK293T cells. The coimmunoprecipitation assays showed that MDK overexpression decreased the binding of STRAD and Mo25a to LKB1; however, it did not affect the LKB1-AMPK interaction (Figs 4C and EV3B). Moreover, gradually increasing MDK in the HEK293T cells was accompanied by gradually decreased LKB1-STRAD-Mo25a association, although the LKB1-AMPKα interaction was not affected (Fig 4D). In contrast, gradually increasing MDK expression affected neither the association of AMPKα with the regulatory β and γ subunits nor the LKB1-AMPKα interaction (Figs 4E and EV3C). To further evaluate the impact of secretion on MDK function, we deleted the three amino acids from 20 to 22 in MDK signal peptide, and named MDK-Del. MDK-Del showed a strong defect in the cleavage of signal peptide and could not detected in the medium (Fig EV3D). Similar to the wild type MDK, MDK-Del interact with LKB1 in cells and attenuated the interaction of LKB1 to STRAD and Mo25 (Fig EV3E and F). Taken together with the association of MDK with LKB1 and STRAD, these results suggest a mechanism by which MDK binds to LKB1 and STRAD and inhibits the formation of the LKB1-STRAD-Mo25a complex, leading to a decrease in LKB1 activity and AMPKα phosphorylation.

**Figure 4.**
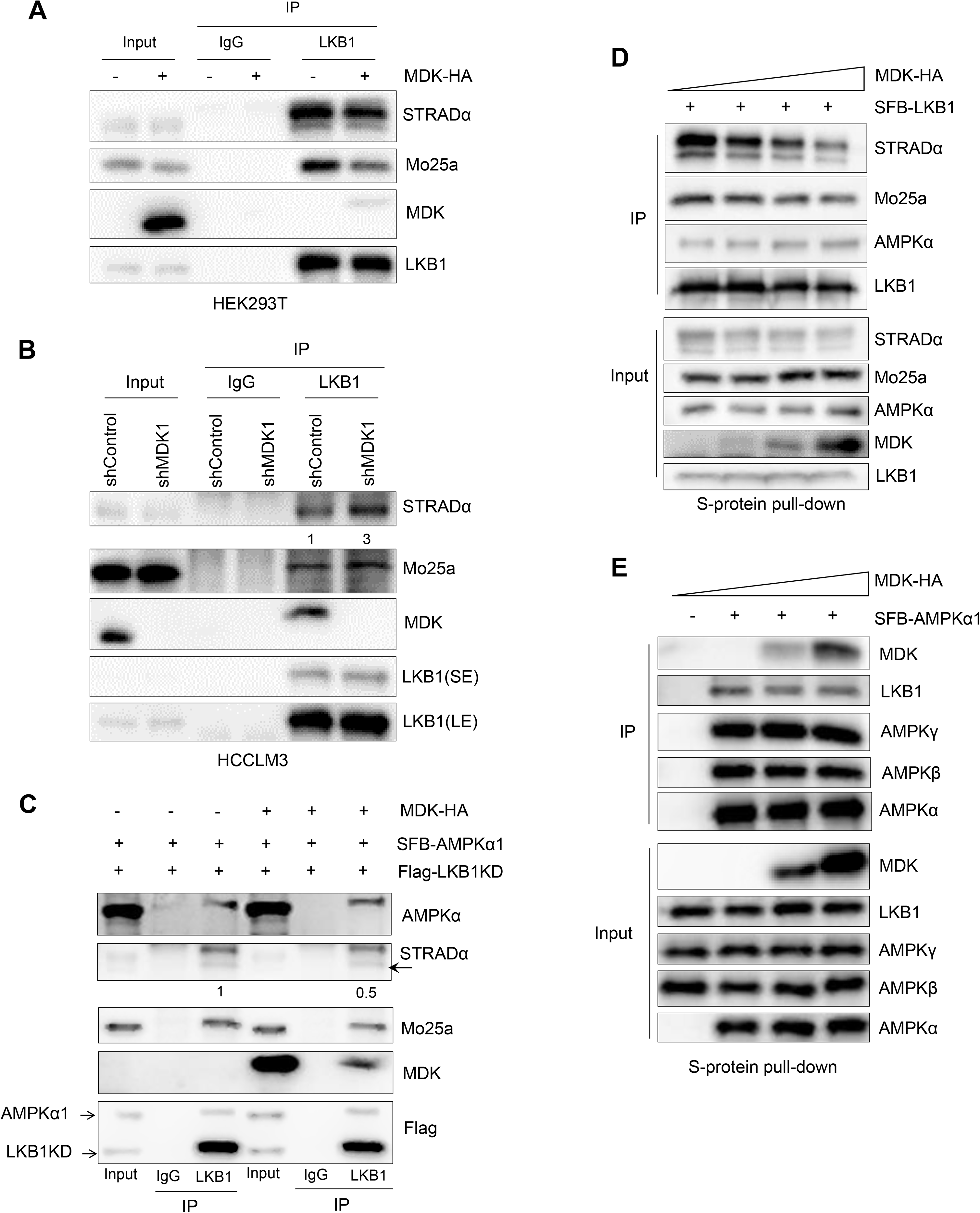
Midkine depolymerized LKB1-STRAD-Mo25 complex. **A and B.** LKB1 was immunoprecipitated from the HEK293T cells transduced with MDK-HA (A) and the HCCLM3 cells transduced with MDK shRNA1 (B), and then, Western blot analysis was performed with antibodies against STRADα, Mo25a, MDK and LKB1. **C.** LKB1 was immunoprecipitated from the HEK293T cells transduced with MDK-HA and the control cells followed by Western blotting with antibodies against AMPKα, STRADα, Mo25a, HA and FLAG. The indicated constructs were expressed in the HEK293T cells for 48 hours, and the cell lysates were subjected to coimmunoprecipitation with primary LKB1 and IgG antibodies. **D and E.** HEK293T cells were cotransfected with different doses of MDK-HA and SFB-tagged LKB1 (D) or SFB-tagged AMPKα1 (E), pulled down with S protein beads and subjected to Western blot analysis with antibodies against STRADα, Mo25a, LKB1, MDK and AMPK subunits.

### Midkine expression is upregulated in cancers

To explore the expression of growth factors in different cancers, we analyzed the expression levels of well-studied growth factors in The Cancer Genome Atlas (TCGA) database. Among these proteins, MDK showed the highest upregulated expression in different types of cancers, even higher than that of established growth factors such as VGF, EGF and TGFB1 (Figs 5A and B, and EV4A), indicating important roles for MDK in cancer. Indeed, cancer was the disease most frequently related to MDK in the disease enrichment analysis (Fig EV4B).

**Figure 5.**
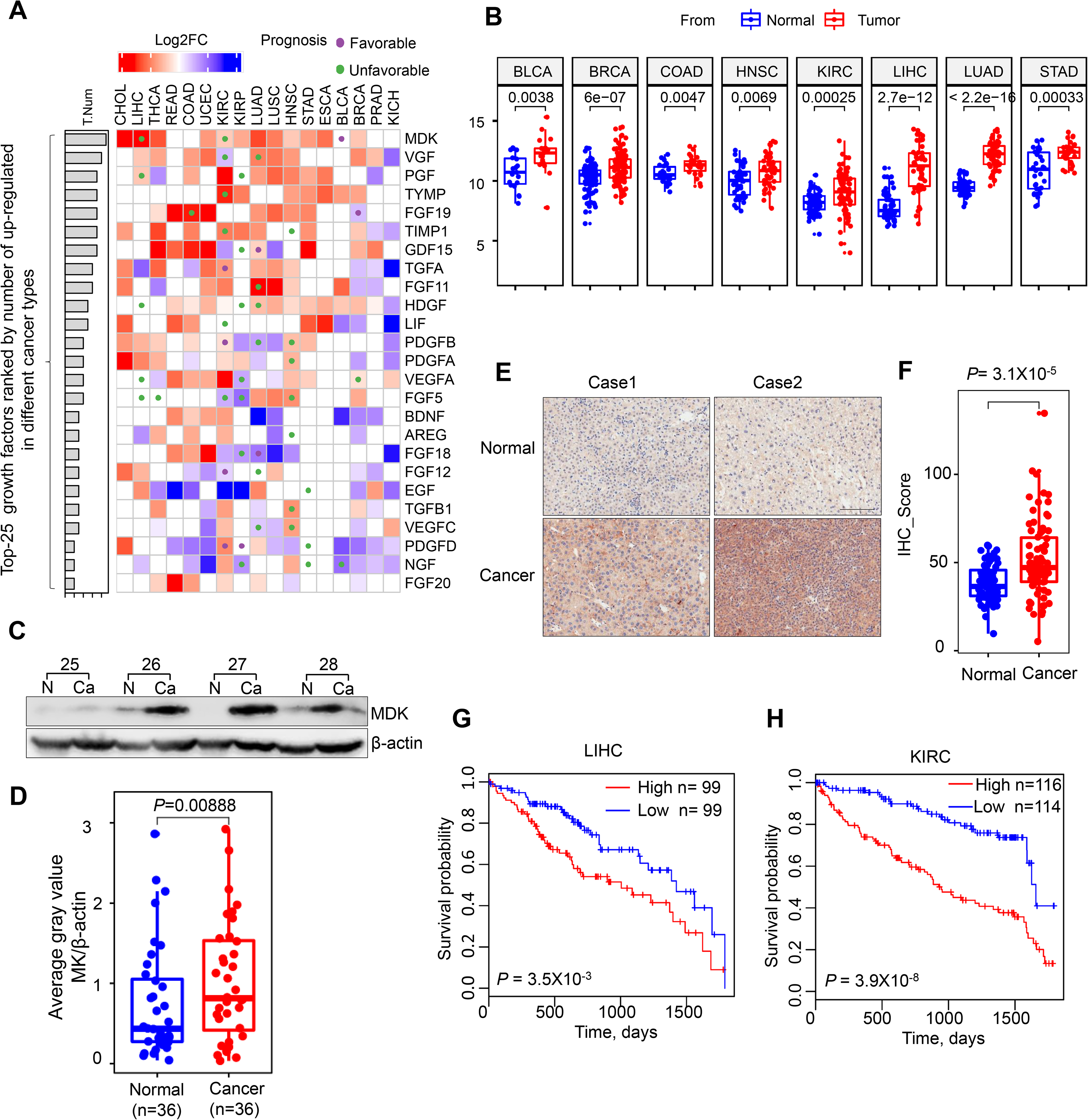
Midkine expression is upregulated in cancer. **A.** Pan-cancer evaluation of the expression and prognostic impact of growth factors. The color of each rectangle represents the log2 transformed fold change (Log2FC) of the mRNA expression for the corresponding growth factor between tumor and normal tissues, and the white rectangles indicate either a Log2FC value equal 0 or differences between tumor and normal tissues that are not significant (linear model approach of limma, P > 0.01). The purple and green circles represent high gene expression correlated with good and poor prognosis, respectively (log rank test, P <0.01). The left bars represent the numbers of cancer types in which a growth factor is upregulated in the tumor tissues compared with the normal tissues (P < 0.01, log2FC > 1). The growth factors are ranked by the numbers, and only the top 25 factors are shown. **B.** Boxplots of the differences in MDK expression in paired normal and tumor tissues of eight types of cancers. The centers of the boxes represent the median values. The bottom and top boundaries of the boxes represent the 25th and 75th percentiles, respectively. The whiskers indicate 1.5-fold of the interquartile range. The dots represent points falling outside this range. The paired P-values were calculated based on Wilcox tests. **C and D.** The expression of MDK in 36 pairs of matched adjacent nontumor (NT) and cancer (Ca) tissues as detected by Western blotting (C), and the distribution of MDK expression in both the NT and Ca samples as represented by boxplots with the expression value normalized by ImageJ software (D). **E and F.** Immunohistochemical staining of MDK in representative adjacent nontumor and HCC specimens (E) and boxplots of the distributions of MDK expression status in 75 paired paraffin-embedded tissues (F). Scale bar, 200 μm. **G and H.** Kaplan-Meier survival curves of LIHC (G) and KIRC (H) patients with data stratified by the expression levels obtained from the TCGA database.

To further examine the expression of MDK in practical samples, we collected 36 pairs of liver cancer tissues and adjacent noncancer tissues. MDK expression was significantly upregulated in the cancer tissues compared to its expression in the adjacent normal tissues (Figs 5C and D, and EV4C). In addition, we also examined MDK expression through tissue microarray assay (TMA), which contained 75 pairs of liver cancer and adjacent normal tissue samples, by immunohistochemistry (IHC). Similarly, MDK was highly expressed in the cancer tissues compared to its expression in the adjacent normal tissues (Figs 5E and F, and EV4D). Notably, higher MDK expression correlated with poor prognosis in both the TCGA liver hepatocellular carcinoma (LIHC) and kidney renal clear cell carcinoma (KIRC) cohorts (Fig 5G and H). Taken together, MDK is highly expressed in most cancers, which suggests that MDK plays important functions in cancer progression.

### Midkine promotes cancer cell proliferation, invasion and tumorigenesis

To clarify the function of MDK in tumorigenesis, we performed both loss-of-function and gain-of-function analyses of MDK in different cancer cell lines. First, we determined the expression level of MDK in a panel of human cancer cell lines. MDK was highly expressed in most cancer cell lines compared to its expression in immortalized normal human liver cells and mammary epithelial cells; however, in some cancer cell lines, MDK expression was almost negligible (Fig EV5A and B). This negative expression of MDK may be due to gene methylation in the *MDK* genomic region or transcriptional regulation in particular cancer cells instead of genome deletion (Fig EV5C).

Considering these expression assessment results, we generated MDK-transduced and short hairpin RNA (shRNA) knockdown cell lines (Figs 6A, E and G and EV5D). Two independent MDK shRNAs both decreased the proliferation of HCCLM3 and HepG2 cells (Figs 6B and F, and EV5 E). In contrast, transducing MDK expression vector in MHCC97H and Bel-7402 cells increased their proliferation (Figs 6C and D, and EV5F). In addition, restoring MDK expression in MDK-knockdown HCCLM3 and HepG2 cells recovered their proliferation ability (Figs 6E and E, and EV5D and E). We also examined the effect of MDK on cell motility and anchorage-independent growth. Knocking down MDK expression significantly decreased the invasion ability of BT549 cells (Fig 6G), and the overexpression of MDK increased the colony-forming ability of Bel-7402 cells in soft agar (Fig 6H). However, MDK-overexpressing cells did not affect wound healing migratory ability (Fig EV5G). To explore the function of MDK in tumor growth *in vivo*, we subcutaneously injected MDK-knockdown or reconstituted HCCLM3 cells and control cells into nude mice. Mice with MDK shRNA-expressing cancer cells produced smaller tumor, measured by volume, and lighter tumor, measured by weight, throughout the experiment than mice transplanted with the control shRNA-infected cells or MDK-reconstituted HCCLM3 cells (Fig 6I-K). In contrast, MDK-overexpressing MHCC97H cells accelerated tumor growth and tumor weight *in vivo* (Fig EV5H and I). Furthermore, the tumors formed by MHCC97H cells overexpressing MDK exhibited downregulated AMPKα phosphorylation compared with the tumors formed by control MHCC97H cells (Fig EV5J). Taken together, these results indicate that MDK promotes the proliferation and tumorigenecity of human cancer cells; however, the mechanism by which intracellular MDK derives tumorigenesis remains unclear.

**Figure 6.**
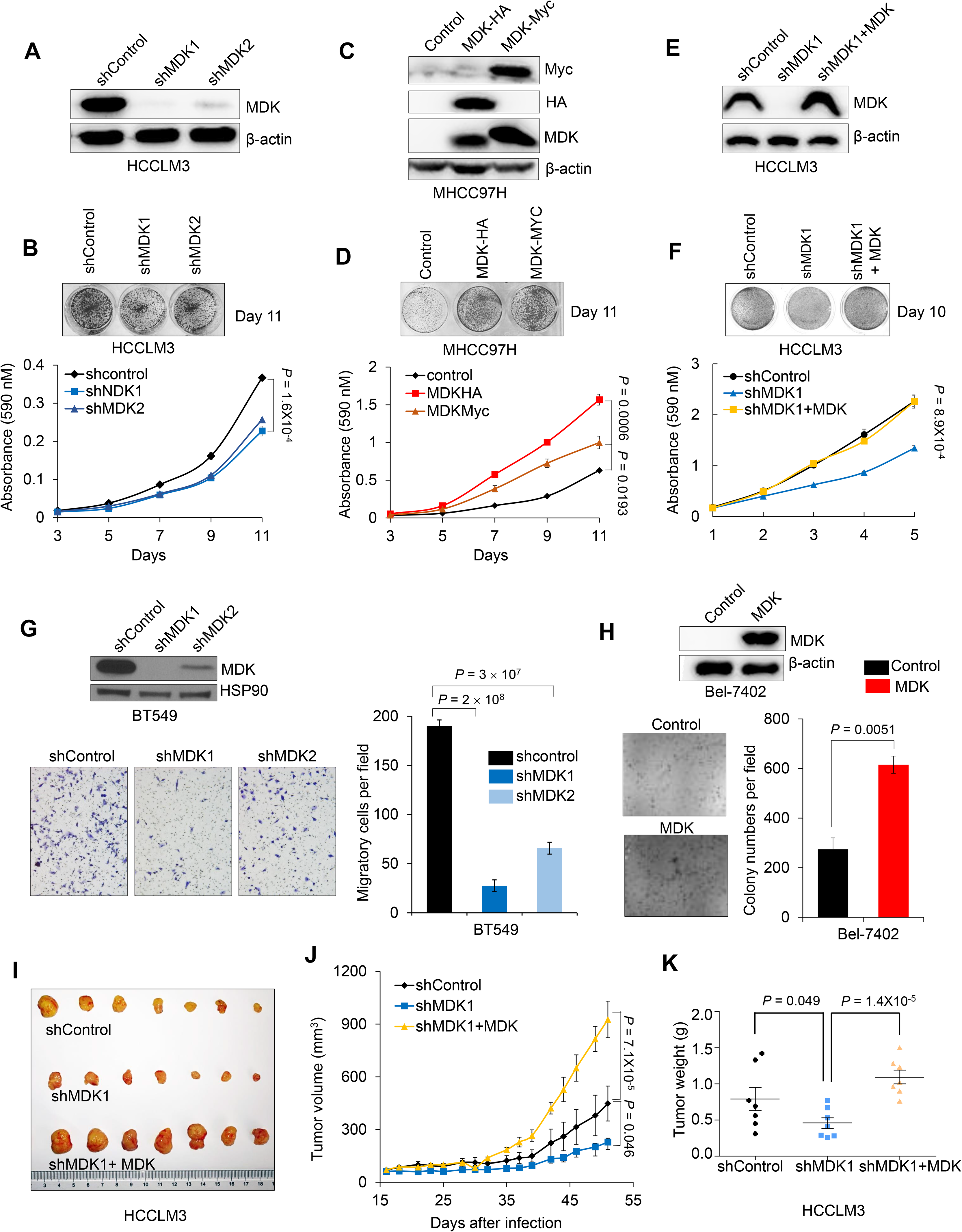
Midkine promotes cancer cell proliferation, invasion and tumorigenesis. **A and B.** Western blotting of MDK and β-actin in HCCLM3 cells transduced with two independent MDK shRNAs (A) and representative images and growth curves of the HCCLM3 cells with MDK knocked down (B). **C and D.** Western blot analysis of MDK, Myc, HA and β-actin in the MDK-transduced MHCC97H cells (C) and representative images and cell growth curves of MHCC97H cells overexpressing MDK (D). **E and F.** Western blot analysis of MDK and β-actin in the HCCLM3 cells transduced with MDK shRNA1 and restored MDK in the MDK-knockdown cells (E) and representative images and cell growth curves of the HCCLM3 cells transduced with MDK shRNA1 and restored MDK in the MDK-knockdown cells (F). **G.** Western blotting of MDK and HSP90 in the BT549 cells transduced with two independent MDK shRNAs and representative images and invaded cell numbers of BT549 cells with MDK knocked down. *n*=3 wells per group. Scale bar, 200 μm. **H.** Western blotting of MDK and β-actin in the MDK-overexpressing Bel-7402 cells and representative images and clone numbers of the Bel-7402 cells with restored MDK expression. *n*=3 wells per group, Bar=200 μm. **I-K.** Tumor images (I), growth curve (J) and weight (K) after subcutaneously injecting mice with HCCLM3 cells transduced with MDK shRNA or reconstituted MDK in MDK-knockdown cells.

### Midkine promotes cancer progression by negatively regulating AMPK signaling

MDK has been reported to promote tumor progression in a diverse manner, but the definitive mechanism remains unclear. Our results showed that MDK accelerated tumor cell proliferation, invasion and *in vivo* tumorigenesis (Fig 6). AMPK is mainly activated under energy stress conditions and suppresses cell division to reduce energy consumption(Gonzalez, Hall et al., 2020). To investigate the role of AMPK in MDK-modulated cell proliferation, we performed a colony forming assay under normal and low-glucose conditions. As expected, MDK-overexpressing cells showed an accelerated proliferation rate under both conditions, and we also observed that, compared to the effects under normal conditions, the overexpression of MDK significantly promoted cell proliferation under low-glucose conditions ( Fig EV6A and B). However, knocking down MDK did not alter the mitochondrial oxygen consumption rate (OCR) or extracellular acidification rates (ECARs) (Fig EV6C and D), thus indicating that MDK did not affect glycolysis or mitochondrial oxidative phosphorylation.

Furthermore, to determine whether MDK decreases LKB1 activity to suppress AMPK activation and thus contributes to cell proliferation, we expressed LKB1 shRNA and CAMKKβ shRNA in MDK-depleted HCCLM3 cells. Knocking down LKB1, but not CAMKKβ, reversed the inhibitory effects of MDK shRNA on cell proliferation and colony formation (Figs 7A and B, and EV6E-H). In addition, reconstitution of LKB1 in its deficient Hela cells, significantly inhibited the cell proliferation caused by MDK overexpression (Fig EV6I-K).

**Figure 7.**
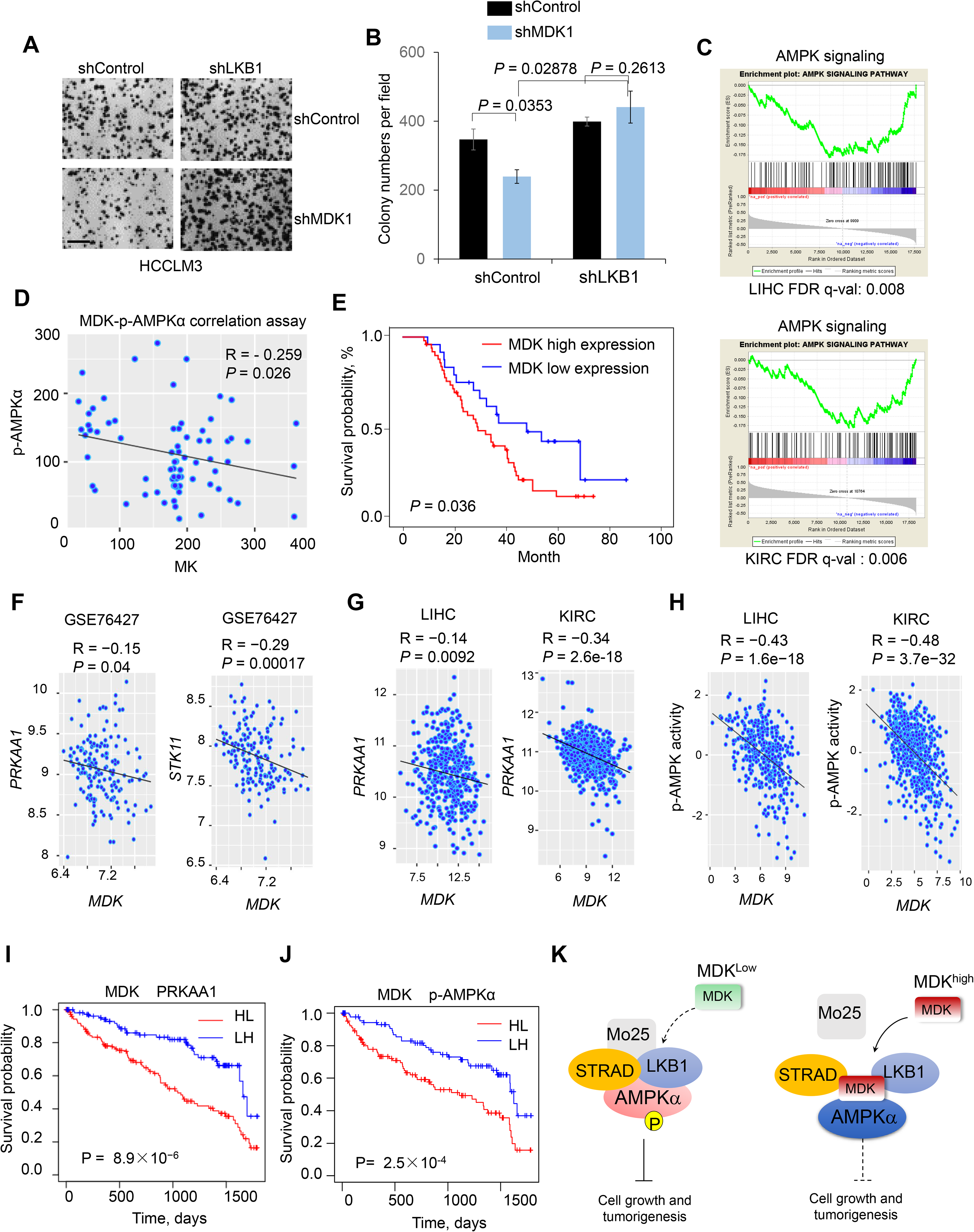
Midkine promotes cancer progression by negatively regulating AMPK signaling. **A and B.** Colony forming assay of the HCCLM3 cells transduced with LKB1 shRNA or in combination with MDK shRNA. Images (A) and quantification (B) of colony formation. *n*=3 wells per group. Scale bar, 200 μm. **C.** GSEA results showing the negative correlations between MDK and the AMPK signaling pathway based on the TCGA LIHC and KIRC cohorts. Genes in the RNA-seq data were ranked by the Pearson coefficients of the correlations between the genes and *MDK*, and the ranked gene list was utilized as the input for the GSEA software program. **D and E.** Scatter plots showing the inverse correlation of MDK with p-AMPKα expression and MDK expression in human hepatocellular carcinoma tumors (D), Kaplan-Meier survival curves of HCC patients with data stratified by MDK expression levels (E) (*n*=74). **F.** Scatter plots showing the inverse correlation of *MDK* with *PRKAA1* (encoding AMPKα1) and *STK11* (encoding LBK1) expression and MDK expression in the GSE76427 data set (*n*=115). **G and H.** Scatter plots showing the inverse correlation of MDK with *PRKAA1* and AMPK in the TCGA LIHC and KIRC cohorts (LIHC n=371, KIRC n=531). Gene expression was obtained from RNA-seq data from the TCGA. AMPK activity was estimated by the expression of their downstream target genes. Statistical significance in (d-h) was determined by Pearson correlation test. R: Pearson correlation coefficient. R, Spearman rank correlation coefficient. **I and J.** Kaplan-Meier survival curves of the TCGA KIRC patients with data stratified by the expression levels of both *MDK* and *PRKAA1* (I) or the expression of *MDK* and the activity of AMPK (J). HL: MDK is high, *PRKAA1 or* AMPK is low; LH: MDK is low, *PRKAA1 or* AMPK is high. Median expression/activity levels were utilized as the thresholds for high and low separation. **K.** Schematic cartoon of the MDK mechanisms of action: high MDK expression depolymerizes the LKB1-STRAD-Mo25 complex and subsequently suppresses the activity of AMPK signaling in human cancers.

To understand whether MDK affects AMPK-related signaling pathways in different cancers, we performed a gene expression correlation-based gene set enrichment analysis (GSEA). The results showed that MDK expression is negatively correlated with the AMPK signaling pathway in several different cancers, such as liver cancer, kidney cancer and breast cancer (Figs 7C and EV7A and B). The IHC analysis of the HCC tissues revealed that MDK protein expression is negatively correlated with p-AMPKα expression (Fig 7D), and high MDK expression correlates with poor prognosis for HCC patients (Fig 7E and Supplementary Table 1).

Next, to investigate the relevance of our findings to human cancer, we analyzed gene expression data from TCGA and Gene Expression Omnibus (GEO) datasets. We found a significant negative correlation between *MDK* and the *PRKAA1* or *STK11* (encoding LKB1) transcript levels in the GSE76427 dataset (Fig 7F). On the other hand, MDK showed strong negative correlations with the expression of *PRKAA1* and the activity of AMPK in the TCGA database (Figs 7G and H, and EV7C and D; see the Methods section for the estimation of AMPK activity). Next, we examined the prognostic value of *MDK* with *PRKAA1* or AMPK using the TCGA dataset of KIRC tumors from 531 patients. The patients with simultaneous high expression levels of *MDK* and low expression levels of *PRKAA1* or low AMPK activity had shorter overall survival in the KIRC cohort (Fig 7I and J). In summary, targeting the high expression of MDK may provide therapeutic benefits in human cancers.

## DISCUSSION

MDK is a growth factor belonging to the pleiotrophin family. This definition, in association with its secretory nature, inspired receptor identification research seeking to elucidate the working mechanism of MDK. Several receptors that bind MDK have been identified: ALK, LRP1, Notch2 and PTPξ (Chen, Bu et al., 2007, Gungor, Zander et al., 2011, Huang, Hoque et al., 2008, Lorente et al., 2011, Muramatsu et al., 2000, Sakaguchi, Muramatsu et al., 2003); however, none of these receptors have high affinity for MDK. MDK was also found to be internalized by cells after secretion, but this point has been largely ignored in previous MDK functional studies. In the present study, we not only confirmed the intracellular transport of MDK (Fig 1B and I-L) but also show the high efficiency of this translocation (Fig EV1A and B). In addition, heparin treatment caused a large decrease in intracellular MDK (Fig 1J-L), indicating that most intracellular MDK was the result of extracellular transport. These results suggested that MDK has highly efficient transportability, which enables it to play important roles in the cytoplasm. However, the detailed internalization mechanism of MDK warrants future investigation.

Indeed, here, we reveal previously undescribed functions of intracellular MDK to modulate LKB1 activity by disrupting the formation of the LKB1-STRAD-Mo25 complex by directly associating with LKB1 and STRAD (Figs 2B-H and 4A-E and EV3A, B , E and F). The LKB1-STRAD-Mo25 complex is the major upstream activator of the energy-sensing AMPK (Hawley et al., 2003). However, LKB1 exhibits weak catalytic activity in cells, and its activation is predominantly stimulated by STRAD and MO25 (Boudeau, Baas et al., 2003, Boudeau, Scott et al., 2004, Hawley et al., 2003, Zeqiraj et al., 2009a). MDK inhibits LKB1 activity by depolymerizing the LKB1-STRAD-Mo25 complex (Fig 2 and Fig 4 ), and its inhibition directly suppressed AMPK activation in cells (Fig 2). Thus, we elucidated a new intracellular function and molecular mechanism of MDK (Fig 7), which will inform us as we further investigate whether this mechanism commonly mediates the activity of other proteins.

The expression of MDK has been reported to be upregulated in diverse types of cancer (Filippou, Karagiannis et al., 2020), which was confirmed in our study (Figs 5A-F and EV4). The elevated expression of MDK was accompanied by decreased patient survivals (Figs 5G and H, and 7I and J), suggesting that MDK serves as a prognostic marker. It has been reported that MDK promotes cancer progression by regulating diverse processes, including cell proliferation, invasion, migration and apoptosis (Dai, Wang et al., 2019, Jono & Ando, 2010, Olmeda, Cerezo-Wallis et al., 2017, Takei, Kadomatsu et al., 2001, Yin, Luo et al., 2002). Consistent with these multiple functions, MDK participates in diverse cellular signaling pathways (Kadomatsu et al., 2013), such as the AKT, ERK and Notch2/JAK pathways(Kishida et al., 2013, Lopez-Valero, Davila et al., 2020, Sandra et al., 2004, Stoica, Kuo et al., 2002). In this study, we found that MDK suppresses the activation of AMPK (Figs 1D-I and 3A-F), which is usually regarded as a tumor suppressor. It will be interesting to know whether MDK contributes to cancer progression through AMPK signaling or other pathways, such as by activating AKT, which we discovered (Figs 1I and J, and EV1E). MDK increases cell proliferation and colony formation under normal and low-glucose culture conditions (Figs 6A-F and H and EV5D-F, EV6A and B), even though MDK did not affect glycolysis or mitochondrial oxidative phosphorylation (Fig EV 6C and D). Therefore, our study proved that the inhibition of AMPK activity by MDK may occur only through LKB1 activity in cancer cells.

AMPK, as a master energy sensor, is activated under energy stress conditions to modulate a series of cell behaviors for maintaining survival, including the repression of cell proliferation. Although AMPK is usually considered to be a tumor suppressor, several studies have also reported that AMPK promotes tumor progression by protecting tumor cells under energy stress conditions (Shaw, 2015). These findings indicate that some individual cases may manifest specific regulatory mechanisms. Here, we also found that the expression of MDK was upregulated overall, but some individuals showed the opposite expression trend (Figs 5B-F, and EV4C and D). It will be interesting to determine the AMPK phosphorylation level and tumor development stage in low-MDK tumors. MDK has been widely presumed to be a diagnostic marker of several different cancers, but the ambiguous regulatory molecular mechanism has postponed its utilization. Here, we elucidate the molecular mechanism by which MDK modulates tumor progression, including the suppression of AMPK signaling, to provide more clues to advance the clinical application of MDK in cancer diagnosis and prognosis.

## ACKNOWLEDGMENTS

We thank members of the Dr. Piao laboratory for helpful discussion. This study is supported by National Natural Science Foundation of China grants (No.81972625, No. 81672440, No.81602155, No.21907093), Dalian Science and Technology Innovation Funding (2019J12SN52), Innovation program of science and research from the DICP, CAS (DICP ZZBS201803).

## AUTHOR CONTRIBUTIONS

H-l.P. and T.X, conceived the project, H-l.P. supervised the project. T.X. and H-l.P. the designed project and T.X performed most of experiments, D.C. performed computational data analysis, T.X., D.C. and H-l.P. analyzed data. X.L, N.Z., W.W., H.C., T.L., R.Y., W.O., H.Q., J.K., C.Z., and S.L. provided significant intellectual input. T.X. and H-l.P. wrote the manuscript with input from all other authors.

---

**Expanded View Fig 1.**
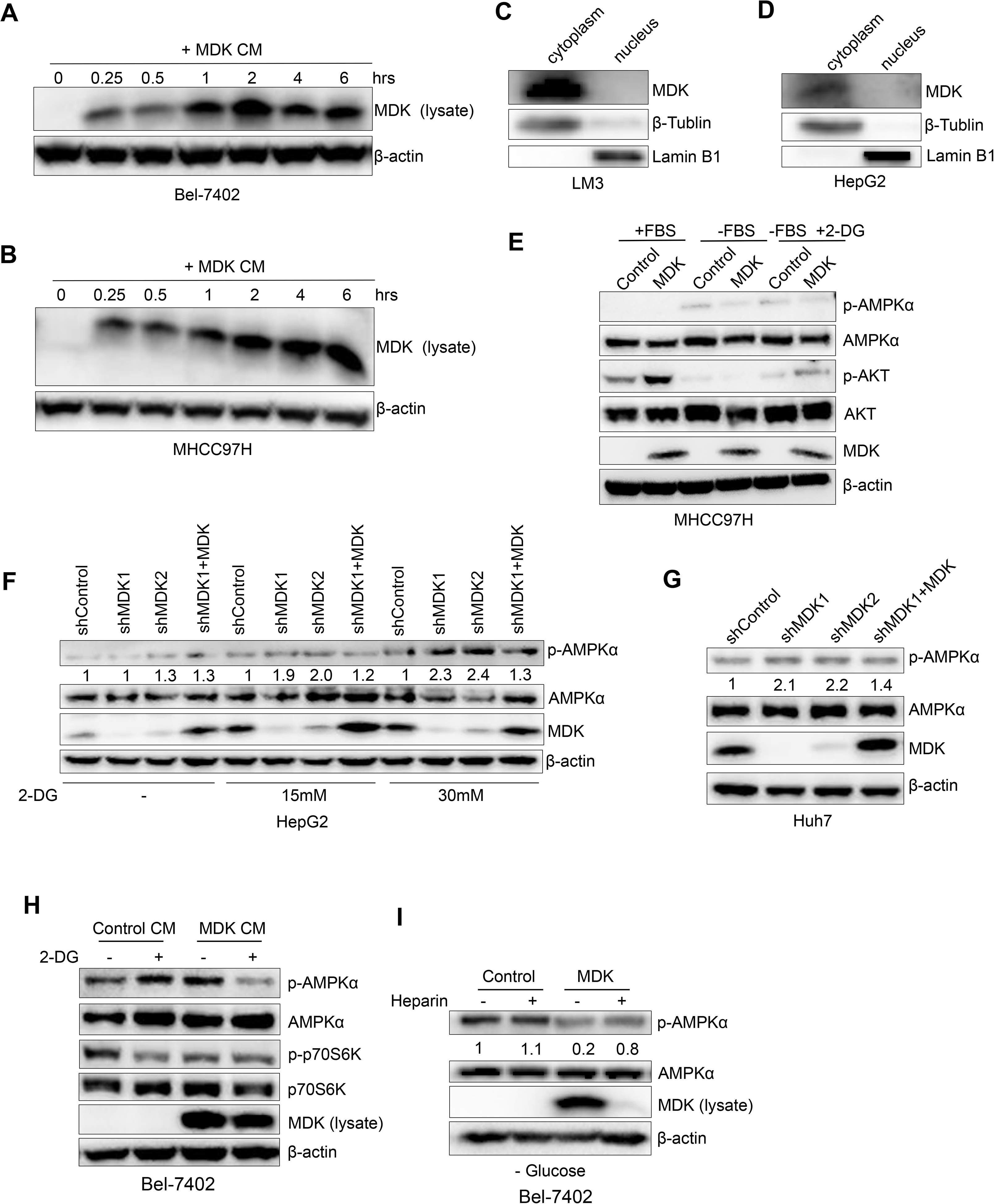

**Expanded View Fig 2.**
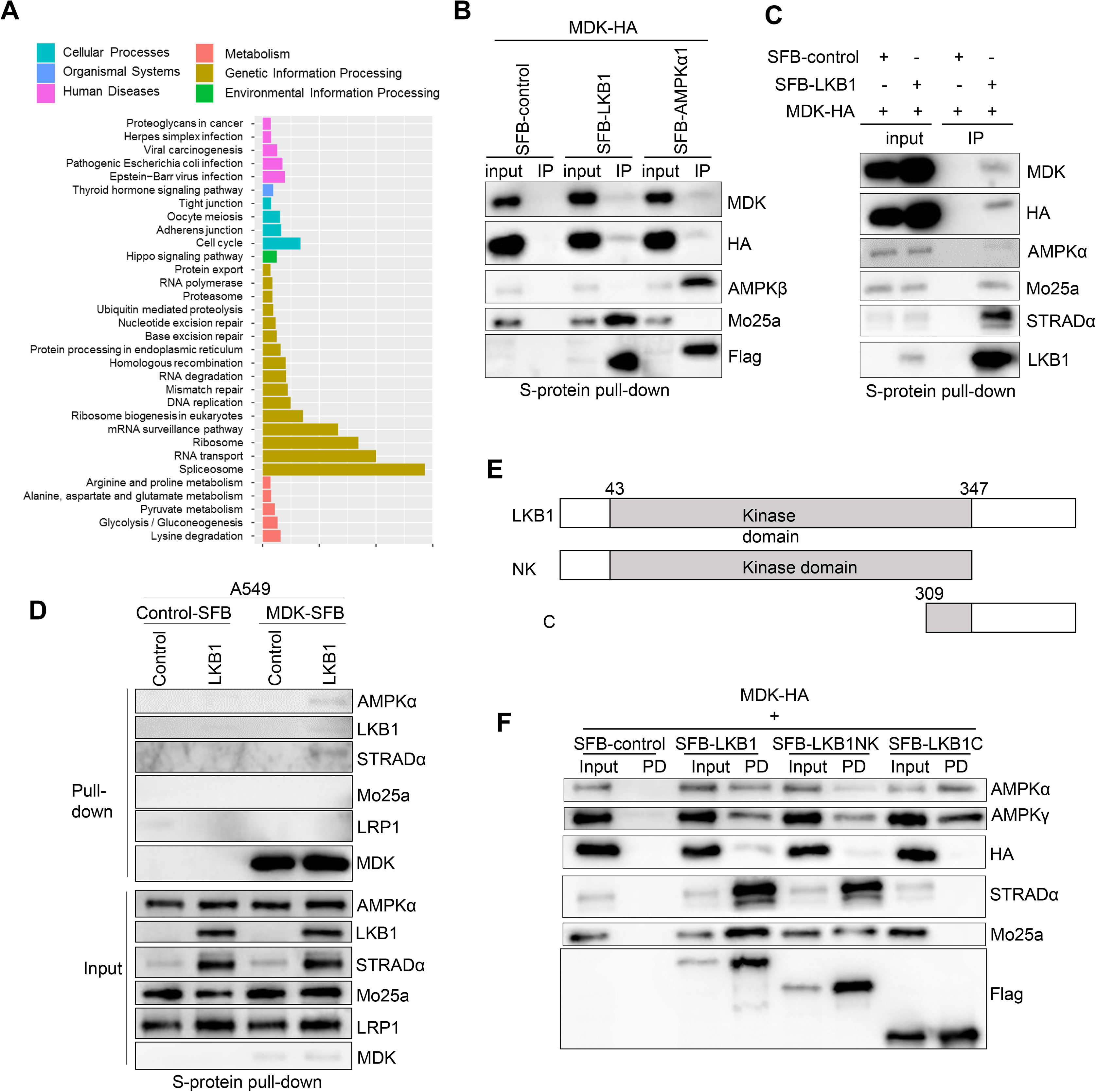

**Expanded View Fig 3.**
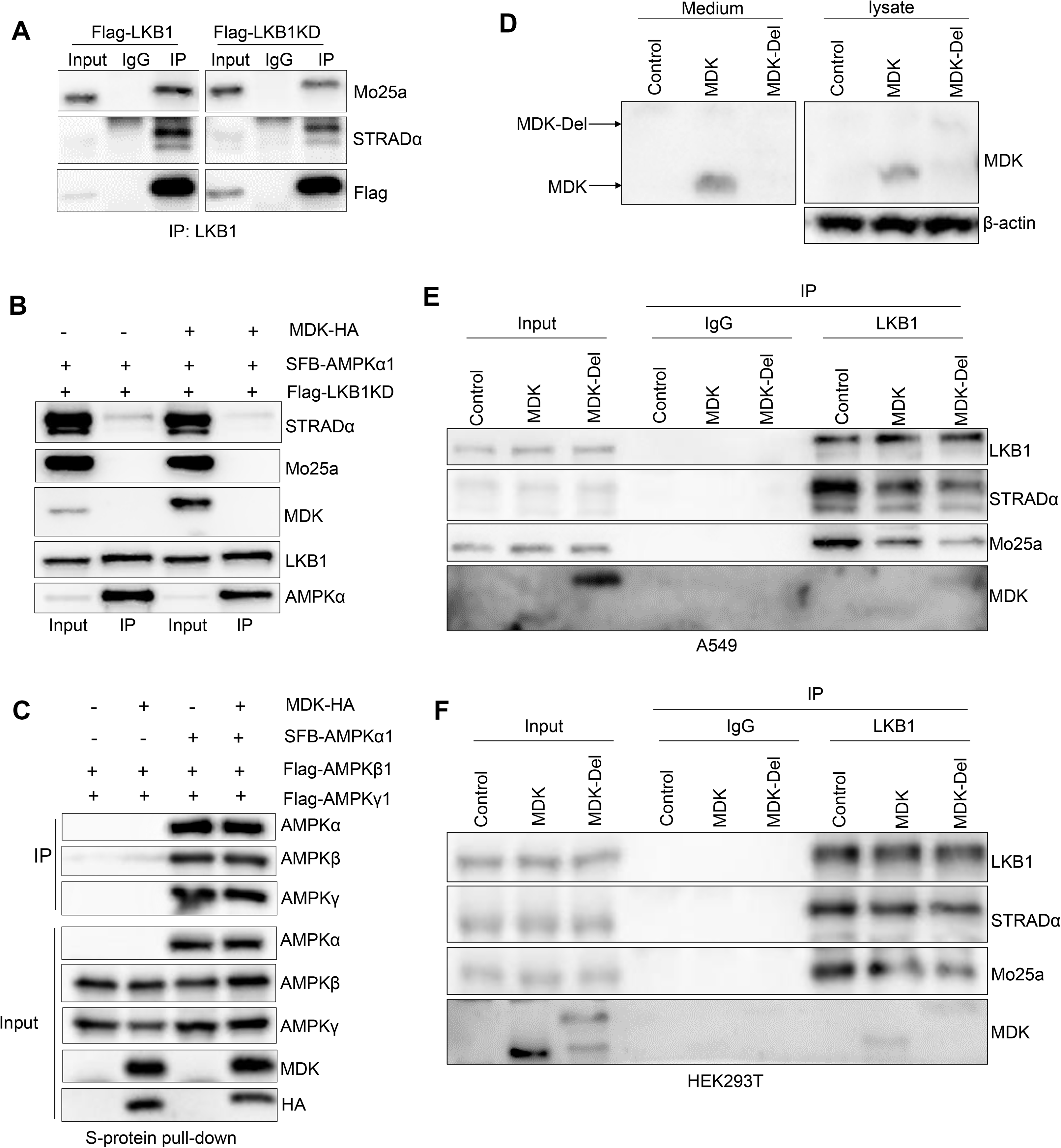

**Expanded View Fig 4.**
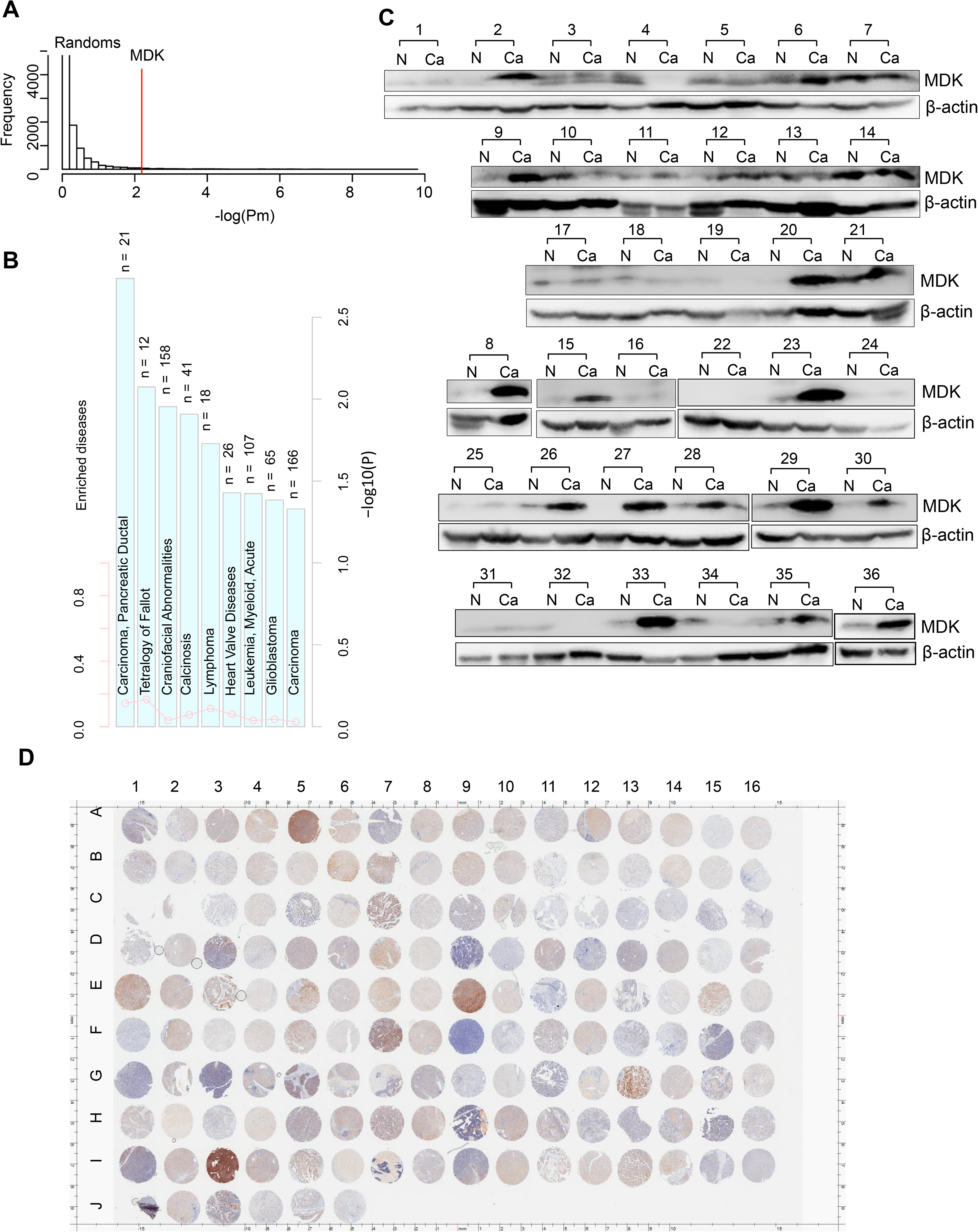

**Expanded View Fig 5.**
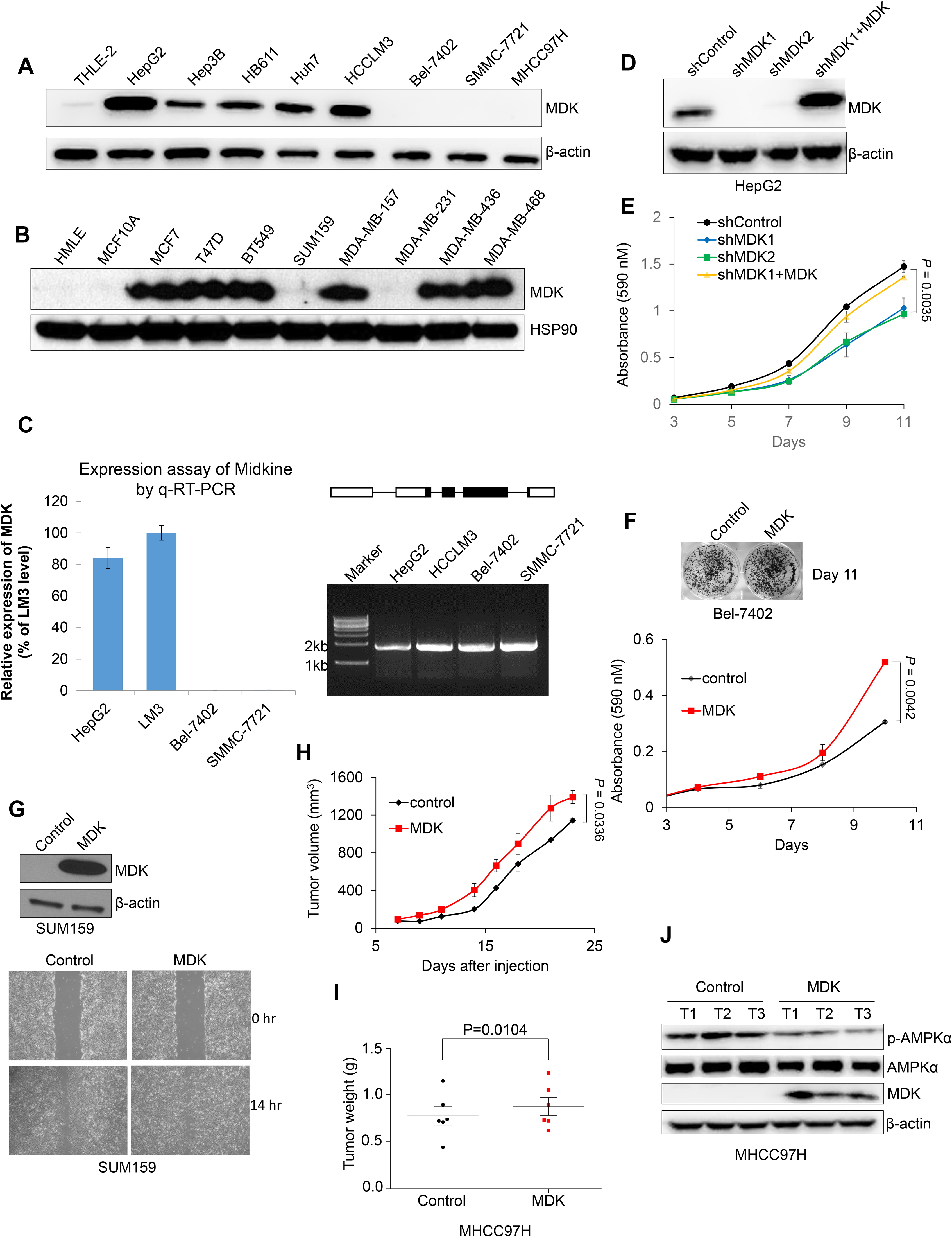

**Supplementary Fig 6.**
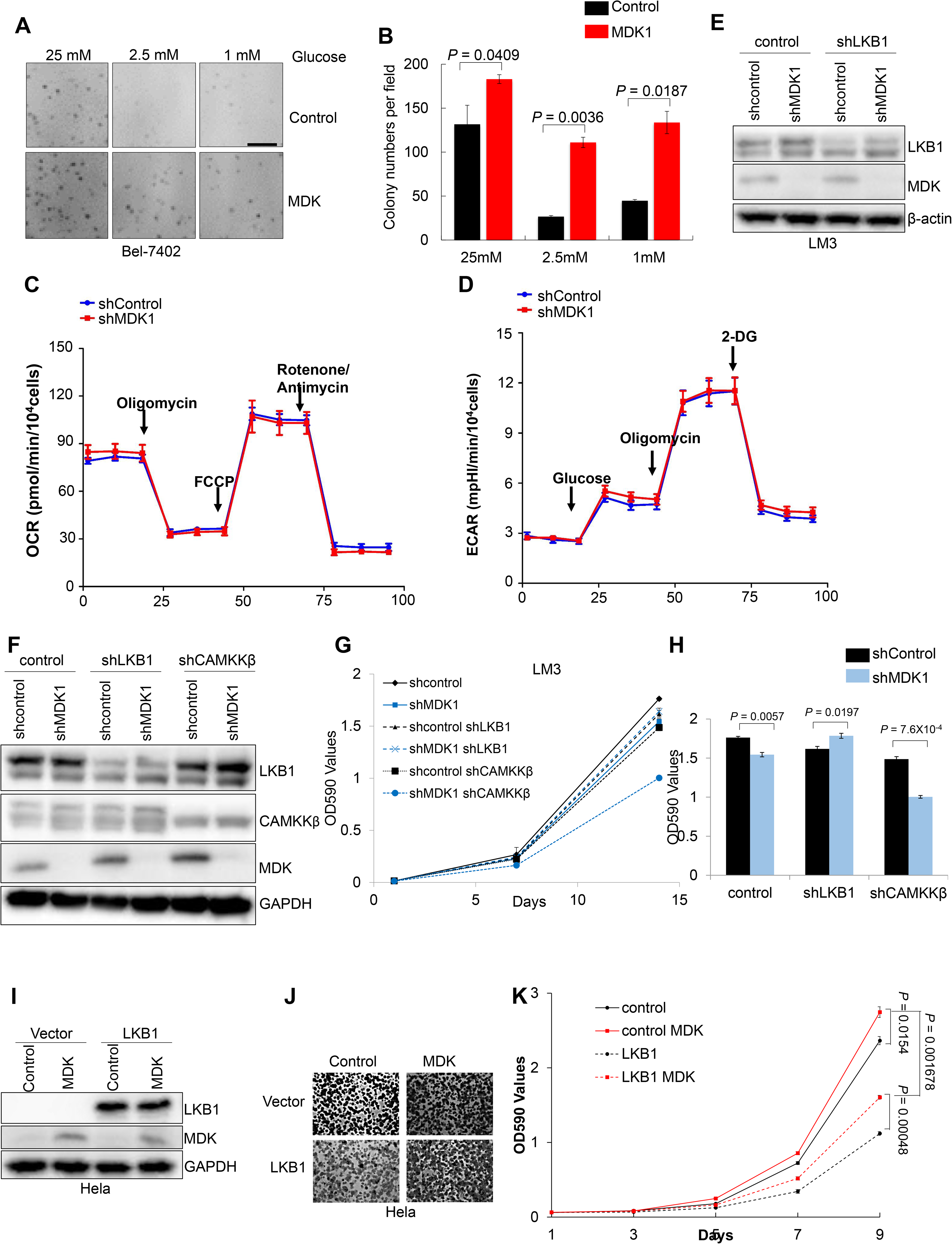

**Supplementary Fig 7.**
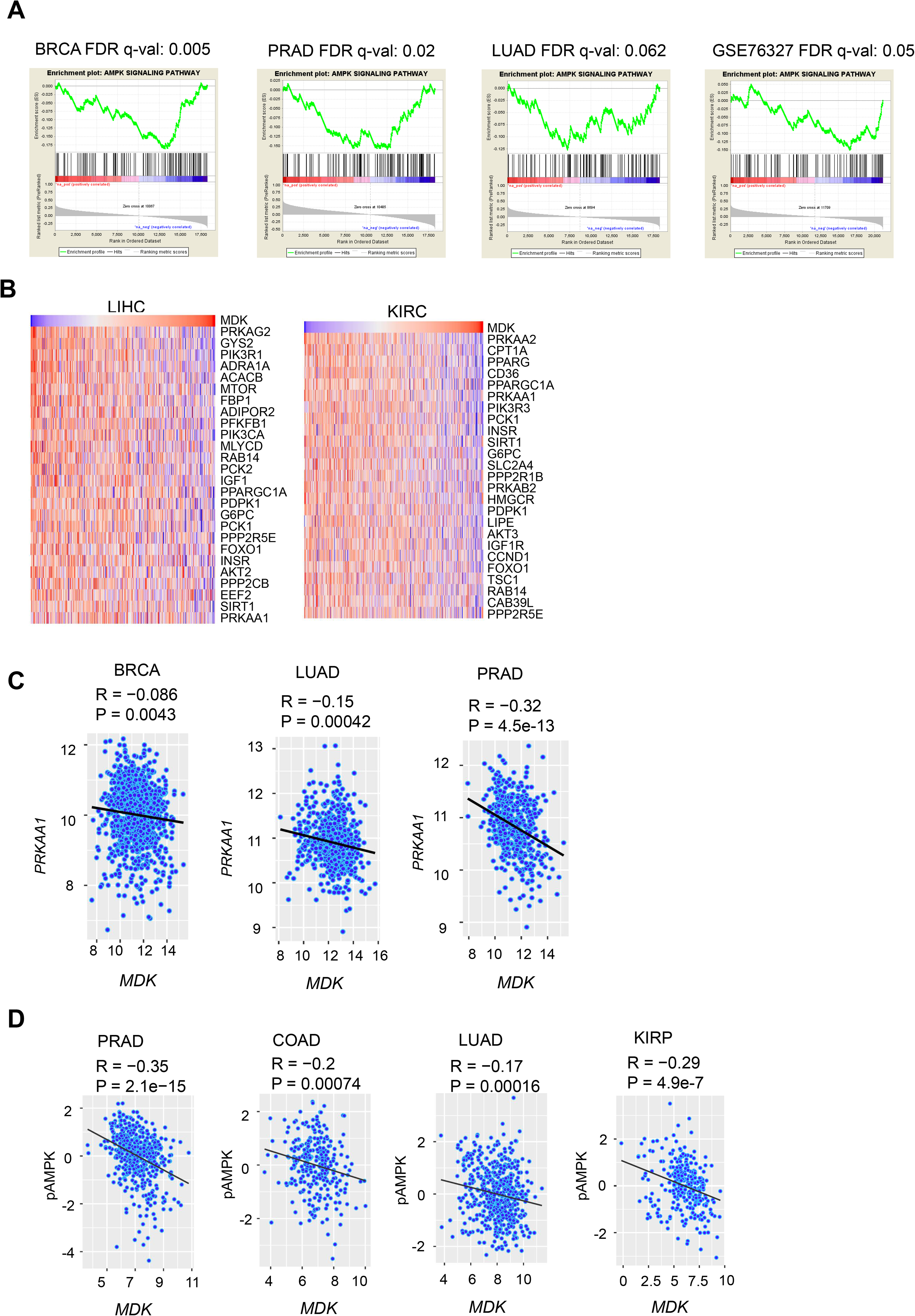

